# Biological and technical variability in mouse microbiome analysis and implications for sample size determination

**DOI:** 10.1101/2024.06.18.599593

**Authors:** Zachary McAdams, Kevin Gustafson, Aaron Ericsson

## Abstract

**Background:** The gut microbiome (GM) affects host growth and development, behavior, and disease susceptibility. Biomedical research investigating the mechanisms by which the GM influences host phenotypes often involves collecting single fecal samples from laboratory mice. Many environmental factors can affect the composition of the GM in mice and while efforts are made to minimize these sources of variation, biological variation at the cage or individual mouse level and technical variation from 16S rRNA library preparation exist and may influence microbiome outcomes. Here we employed a hierarchical fecal sampling strategy to 1) quantify the effect size of biological and technical variation and 2) provide practical guidance for the development of microbiome studies involving laboratory mice.

**Results:** We found that while biological and technical sources of variation contribute significant variability to microbiome alpha and beta diversity outcomes but their effect size is 3- to 30-times lower than that of the experimental variable in the context of an experimental group with high intergroup variability. After quantifying variability of alpha diversity metrics at the technical and biological levels, we then simulated whether sequencing multiple fecal samples from individual mice could improve effect size in a two-group experimental design. Collecting five fecal samples per mouse increased effect size achieving the maximum 5% reduction in the required number of animals per group. While reducing the number of animals required, sequencing costs were dramatically increased.

**Conclusions:** Our data suggest that the effect size of biological and technical factors may contribute appreciable variability to an experimental paradigm with relatively low mean differences. Additionally, repeated sampling improves statistical power however, its application is likely impractical given the increased sequencing costs.

## Background

The gut microbiome (GM) is a diverse community of microorganisms colonizing the gastrointestinal tract that supports crucial roles in influencing host health and disease through the production of microbial metabolites [1,2], modulation of host immune responses [3], and more. Many of these microbiome-mediated mechanisms have been revealed by leveraging germ-free animals[1], standardized complex communities [4,5], and more recently, synthetic microbial communities [6,7]. The use of laboratory mice in microbiome science has dramatically advanced the field as microbiome experimental paradigms may be applied to wild-type or disease-specific mouse models in a relatively quick and affordable manner. However, variability in the laboratory mouse gut microbiome should be considered when designing microbiome experiments as many environmental factors (e.g., diet, housing, vendor of origin) affect the composition of the gut microbes and, in turn, host phenotypes [8,9].

The principles of the 3Rs (replacement, reduction, and refinement) have served as a guide for the humane use of animal models in biomedical research since first being published in 1959 [10]. Progress to replace animal models in microbiome research has been made as evidenced by the development of gut-like bioreactors [11]. These systems do not truly replicate host physiology and lack the ability to measure host phenotypes including behavior and disease outcomes. Additionally, given that fecal sampling is often a non-invasive sampling procedure, there is little need to refine the methods involved in characterizing the GM when examining fecal contents. There is an opportunity to reduce animal numbers by minimizing the biological and technical variability encountered when characterizing the gut microbiome [12]. Efforts to limit this variability typically involve eliminating complicating experimental variables [9], reduction in animal housing density [13], and standardizing sample processing and analysis [14]. However, practical approaches to minimize the inherit variability of an individual subject, or individual samples for that matter, have not yet been explored.

Next-generation sequencing (NGS) techniques are often used to characterize microbial communities given their ability to identify both culturable and unculturable microbes with high taxonomic resolution. Specifically, targeted amplicon sequencing is a NGS technique that involves the amplification of a specific marker gene – typically the 16S rRNA gene for bacterial communities [15,16] – before sequencing those amplicon libraries. This approach is relatively cost-effective and requires minimal computational resources to characterize microbial communities compared to untargeted, metagenomic shotgun sequencing [16]. In 16S rRNA amplicon sequencing, there are many points along the microbial community analysis pipeline at which variability may be introduced including sample collection and processing [14,17,18], PCR during library preparation [19,20], and bioinformatic algorithms [14]. Evaluating the contribution of inherent biological and technical variability in microbiome experiments is important as it allows for the development of strategies to minimize that variability, ultimately reducing the minimum number of animals required in biomedical research.

To quantify the variability and provide practical guidance for the development of microbiome studies using laboratory mice, we leveraged a multi-level hierarchal fecal sampling strategy of two mouse colonies housed at our institution which are colonized with GMs originating from The Jackson Laboratory or Envigo. The gut microbial communities harbored within these two mouse colonies exhibit robust differences in community richness, diversity, and taxonomic composition [4,21,22]. Additionally, these microbial communities are known to affect many host phenotypes including growth and development, behavior, and disease susceptibility, [4,5,23–25]. In applying a hierarchal sampling strategy to these two standardized, complex GMs, we quantified biological and technical variability observed in the context of an experimental factor with well-defined intergroup variability to 1) quantify the effect size of biological and technical variables on microbiome outcome measures, 2) compare the effect size of these factors to that of the experimental variable, and 3) provide simulated sample size calculations and cost analysis demonstrating practical outcomes of repeatedly sampling individual subjects for 16S rRNA amplicon sequencing in microbiome studies.

## Methods

### Ethics Statement

This study was conducted in accordance with the recommendations set forth by the Guide for the Care and Use of Laboratory Animals and was approved by the University of Missouri institutional Animal Care and Use Committee (MU IACUC protocol #36781).

### Mice

C57BL/6J (B6) mice (Jackson Laboratory, Bar Harbor, ME, USA) were set up as breeding trios. The pups from these trios were cross-fostered onto CD-1 dams (Mutant Mouse Resource and Research Center [MMRRC], University of Missouri, Columbia, MO, USA) harboring either GM_Low_ (The Jackson Laboratory-origin) or GM_High_ (Envigo-origin) within 24 hours of birth to transfer the GMs to the surrogate pups. The surrogate pups generated from these cross-fostered litters were confirmed to have been colonized with the GM of their respective CD-1 donor surrogate dam via 16S rRNA gene amplicon sequencing and were used as colony founders. The mice used in this study were the 7th generation of these colonies. GM_Low_ and GM_High_ CD-1 mice were originally generated from C57BL/6J (Jackson Laboratory, Bar Harbor, ME, USA), or C57BL/6NHsd (Envigo, Indianapolis, IN, USA) surrogate dams via surgical embryo transfer^28^.

All C57BL/6J mice used in this study were housed two animals per cage under barrier conditions in microisolator cages (Thoren, Hazleton, PA, USA) with aspen chip bedding. Mice had *ad libitum* access to irradiated LabDiet 5053 chow (LabDiet, St. Louis, MO, USA), and autoclaved tap water. The facility maintains all animals under 12:12 light/dark cycle. Mice were found to be free of *Bordetella bronchiseptica; Filobacterium rodentium; Citrobacter rodentium; Clostridium piliforme Corynebacterium bovis*; *Corynebacterium kutscheri*; *Helicobacter* spp.; *Mycoplasma* spp.; *Pasteurella pneumotropica*; *Pneumocystis carinii*; *Salmonella* spp.; *Streptobacillus moniliformis*; *Streptococcus pneumoniae*; adventitious viruses including H1, Hantaan, KRV, LCMV, MAD1, MHV, MNV, PVM, RCV/SDAV, REO3, RMV, RPV, RTV, and Sendai viruses; intestinal protozoa including *Spironucleus muris*, *Giardia muris*, *Entamoeba muris*, trichomonads, and other intestinal flagellates and amoebae; intestinal parasites including pinworms and tapeworms; and external parasites including all species of lice and mites via quarterly sentinel testing.

### Fecal Collection

Fecal samples were collected from individual mice beginning between 6:30 and 7:00 AM. Briefly, mice were individually placed into clean, empty cages. Mice were allowed to freely defecate up to three pellets within 15 minutes to prevent potential temporal variation between samples. Pellets were collected with autoclaved toothpicks and individually placed into 2.0 mL round-bottom tubes with a single 0.5 cm steel ball for processing. Samples were stored at −80°C until DNA extraction.

### DNA extraction

DNA was extracted from each sample using a modified QIAamp PowerFecal Pro DNA extraction kits (QIAGEN). Samples were homogenized at for 10 min at 30 Hz using a TissueLyser II (QIAGEN). DNA extraction continued per manufacturer instructions. DNA yields were quantified via fluorometry (Qubit 2.0, Invitrogen, Carlsbad, CA) using quant-iT BR dsDNA reagent kits (Invitrogen). When appropriate, DNA yields were normalized to 3.51 ng/ μL using Buffer C6 (QIAGEN).

### 16S rRNA library preparation and sequencing

Library preparation and sequencing were performed at the University of Missouri Genomics Technology Core. Bacterial 16S rRNA amplicon libraries were constructed in technical triplicates via amplification of the V4 region of the 16S rRNA gene with universal primers (U515F/806R) [33] previously developed against the V4 region, flanked by Illumina standard adapter sequences. PCR was performed as 50 µL reactions containing 100 ng metagenomic DNA, dual-indexed forward and reverse primers (0.2 µM each), dNTPs (200 µM each), and Phusion high-fidelity DNA polymerase (1U, Thermo Fisher). Amplification parameters were 98°C^(3 min)^ + [98°C^(15 sec)^ + 50°C^(30 sec)^ + 72°C^(30 sec)^] × 25 cycles + 72°C^(7 min)^ [21]. Amplicon pools were combined then purified by addition of Axygen Axyprep MagPCR clean-up beads to an equal volume of 50 µL of amplicons and incubated for 15 minutes at room temperature. Products were washed multiple times with 80% ethanol and the pellet was resuspended in 32.5 µL EB buffer (Qiagen), incubated for two minutes at room temperature, and then placed on the magnetic stand for five minutes. The final amplicon pool was evaluated using an Advanced Analytical Fragment Analyzer automated electrophoresis system, quantified using quant-iT HS dsDNA reagent kits, and diluted according to the Illumina standard protocol for sequencing as 2×250 bp paired-end reads on the MiSeq instrument.

### Informatics

All 16S rRNA amplicons were processed using the Quantitative Insights Into Microbial Ecology 2 (QIIME2) framework v2021.8 [34]. Illumina adapters and primers were trimmed from forward and reverse reads with cutadapt [35]. Untrimmed sequences were discarded. The paired-end reads were then truncated to 150 base pairs and denoised into amplicon sequence variants (ASVs) using DADA2 [36]. Paired-end reads were merged based on a minimum overlap of 12 base pairs. Merged sequences were filtered to between 249 and 257 base pairs in length. Unique sequences were then assigned a taxonomic classification using a sklearn algorithm and the QIIME2-provided 99% non-redundant SILVA v138 [37] reference database trimmed to the 515F/806R [33] region of the 16S rRNA gene. The resulting feature table of ASV counts per sample was rarefied to a uniform depth of 39,293 features per sample.

### Statistical Analysis

All statistical analyses were performed using various libraires within R v4.2.2 [38]. Initial effects of GM and sex on univariate data were assessed by fitting data to a linear mixed model then evaluating differences using a two-way analysis of variance (ANOVA). Precision of univariate data was assessed using the coefficient of variance (CV = σ/μ). Main effects of GM and sex on multivariate data were initially assessed using a two-way permutational multivariate analysis of variance (PERMANOVA) with weighted (Bray-Curtis) distances using the *adonis2* function from the *vegan* library [39,40]. Nested ANOVAs (or PERMANOVAs) were then uses to evaluate significant effects of technical and biological variables in both univariate and multivariate data using the following model: *outcome ∼ GM* / *cage* / *mouse* / *replicate*. Nested ANOVAs and PERMANOVAs were performed using the *rstatix* [41] and *vegan* [39,40] libraries, respectively. Effect size was calculated using eta squared (η^2^ = Sum of Squares_Variable_ / Total Sum of Squares).

Dissimilarities at the replicate, mouse, and cage levels were calculated by first creating distance matrices using weighted (Bray-Curtis) or unweighted (Jaccard) dissimilarities. Distance values were then averaged for each sample, mouse, and cage to evaluate precision. Principal coordinate analyses (PCoA) were generated using quarter-root-transformed feature tables. The PCoA was performed using the *ape* [42] library with a Cailliez correction.

Power calculations were performed on a simulated experiment in which there are two groups with an expected mean difference low, medium, and high differences. The expected variance was evaluated based on the following equation with the respective variance values calculated based on the present data: 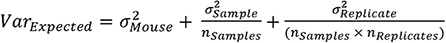[26]. That variance was then used to calculated Cohen’s *d* 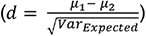 generating medium to high effect sizes. In our simulated experiment, we assumed the number of technical replicates (*n_Replicate_*) to be one and varied the number of samples (*n_Samples_*) collected per mouse. Assuming □ = 0.05 and β = 0.8, we determined the effect size achieved for each mean difference and the minimum number of mice required per group.

## Results

### Supplier-origin microbiomes differ in richness and diversity

To determine the relative effect size of biological and technical variability contributed to 16S rRNA microbial community analysis, we employed the hierarchical study design depicted in **Figure 1A**. We sampled C57BL/6J (B6) mice which harbored either a Jackson Laboratory- or Envigo-origin GM. Four cages of pair-housed mice per sex per GM were sampled (4 cages × 2 sexes × 2 GMs × 2 mice = 32 mice). Three fecal pellets were collected from each mouse (32 mice × 3 pellets = 96 pellets). Fecal DNA extracted from each pellet was then sequenced in triplicate (96 extractions × 3 16S rRNA libraries = 288 libraries). Thus biological variability was assessed at the cage and mouse levels while technical variability of 16S rRNA library preparation was evaluated.

**Figure 1.**
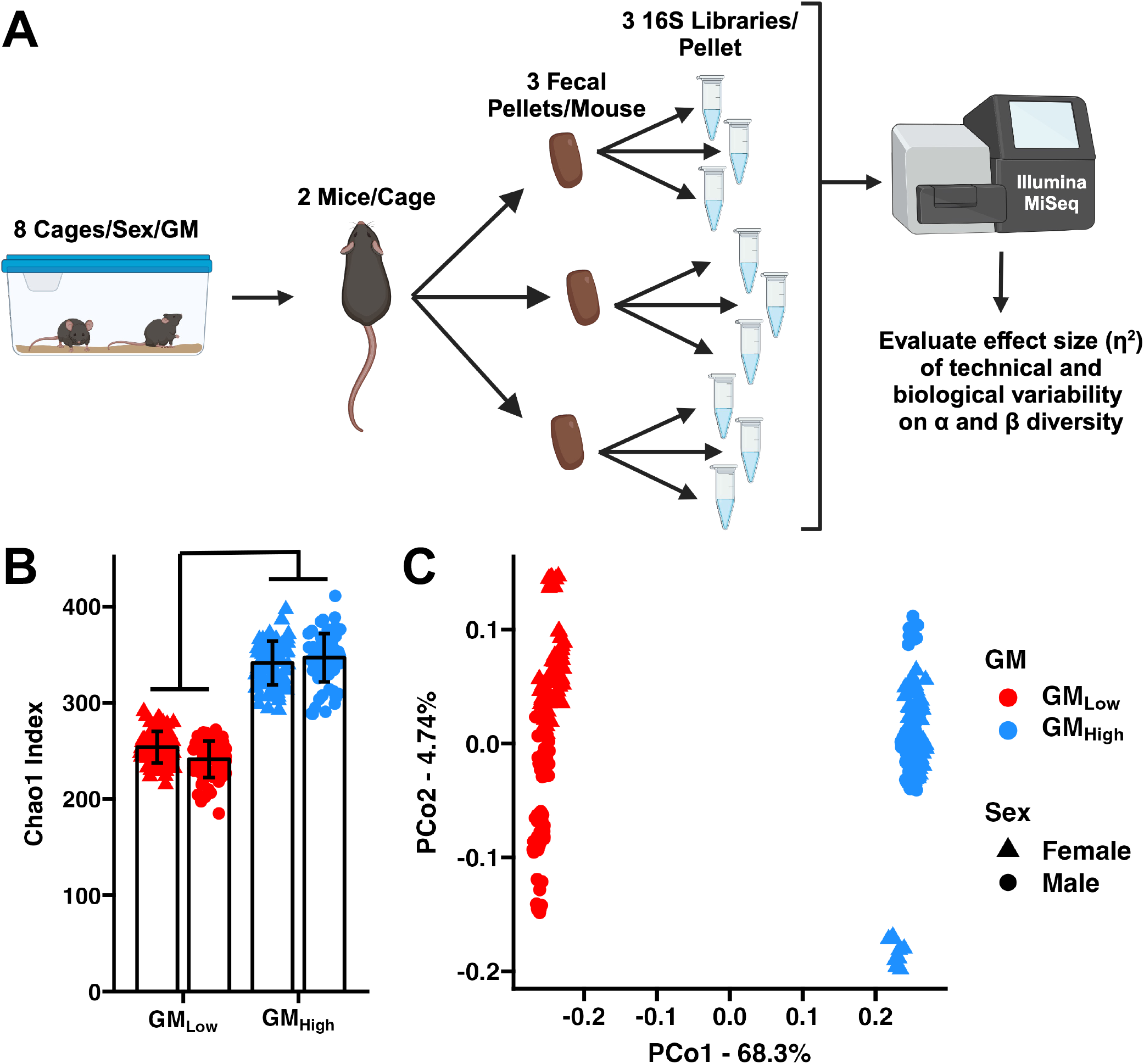
Vendor-origin microbiomes differ in richness and microbial composition. (**A**) Hierarchical study design depicting fecal sampling and sequencing approach from two colonies of mice with distinct vendor-origin microbiomes. (**B**) GM_Low_ and GM_High_ microbiomes differ in richness (F = 241, *p* < 0.001) but not by sex (F = 0.314, *p* = 0.580), two-way ANOVA. (**C**) PCoA depicting vendor-dependent effects on weighted beta diversity, (GM: F = 394, *p* < 0.001, Sex: F = 22.1, *p* < 0.001, GM × Sex: F = 18.9, *p* < 0.001), two-way PERMANOVA.

B6 mice with a Jackson Laboratory-origin GM exhibited a less rich and diverse community compared to mice with an Envigo-origin GM (**Figure 1B**, **Supplementary Figure 1**). Therefore, these GMs were referred to as GM_Low_ and GM_High_, respectively. As no sex-dependent effects were observed on any univariate alpha diversity measurement from either colony; we removed sex as a main effect from our analysis to increase the interpretability of the nested models described below. Assessment of beta diversity using weighted, Bray-Curtis distances revealed significant effects of vendor and sex, however, vendor-dependent effects were the dominant source of separation using a principal coordinate analysis (PCoA, **Figure 1C**). Samples clustered by vendor and separated along the first axis which captured 68.3% of variation in the dataset.

### Low richness microbiome exhibits increased variability in presence-based alpha diversity metrics

We first assessed the precision of alpha diversity measurements at the technical and biological levels by calculating the coefficient of variation (CV) for metrics measuring richness (Observed Richness and Chao1 Index) and diversity (Shannon and Simpson Indices). GM_Low_, the low-richness Jackson Laboratory-origin GM, exhibited a significantly increased CV at the replicate and mouse levels when evaluating richness metrics (**Supplementary Figure 2**). GM_Low_ Simpson index measurements also exhibited an increased CV at the replicate and mouse, but not cage levels. Shannon index measurements for both GMs exhibited similar CV values at each level. These data suggest that presence-based measurements are susceptible to technical and biological variation in low richness GM, ultimately decreasing precision.

### Technical and biological variables contribute minimal effect sizes relative to experimental variable

To evaluate the effect size that the biological and technical variables contributed to microbiome outcome measures, we applied a nested ANOVA model to univariate and multivariate data. In addition to the main effect of GM, technical (16S rRNA library preparation) and biological variation (mouse and cage) contributed significant variation to each evaluated microbiome outcome measure, with the exception of technical replicates not affecting richness-based measurements (Observed Richness and Chao1 Index, **Figure 2**). Upon comparing effect sizes, we found that relative to the main experimental effect of GM (η^2^ = 0.62 ± 0.20), biological variability at the cage (η^2^ = 0.21 ± 0.12) and mouse levels (η^2^ = 0.13 ± 0.086) and technical variability at the replicate level (η^2^ = 0.027 ± 0.014) contributed an effect size three- to twenty-times lower than the experimental variable of interest (**Figure 2**). When evaluating these effects within each GM, large effects of cage and individual mouse were observed, especially in multivariate analyses. The relative effect size at the technical level was low compared with cage and mouse effect sizes (**Supplementary Figure 3**). Collectively, these data suggest that biological and technical variables contribute significant variation to microbiome outcome measures, but this effect is minimal relative to an experimental variable of interest.

**Figure 2.**
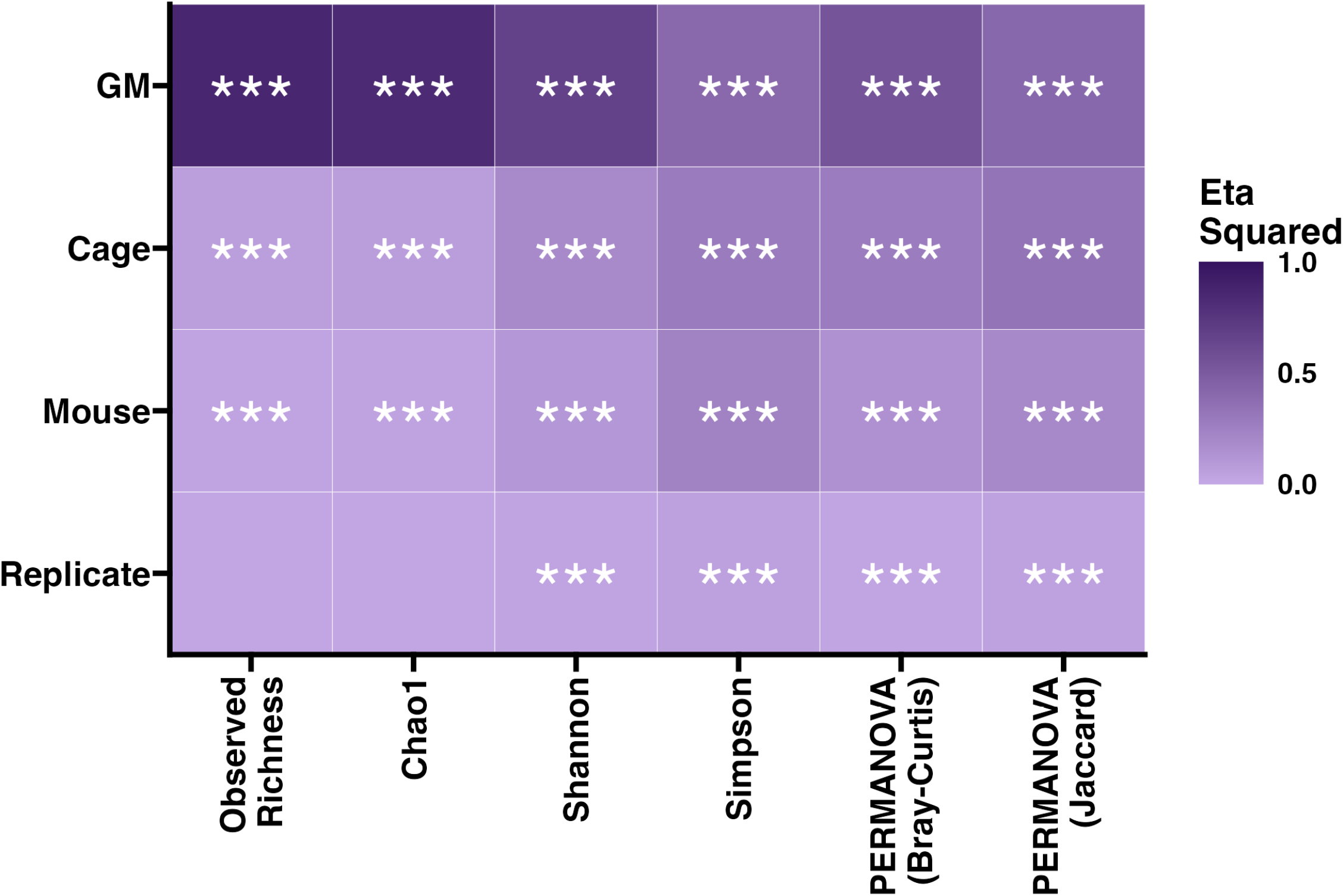
Biological and technical factors contribute small effect sizes relative to experimental variable. Heatmap depicts the relative effect size contributed by technical, biological, and experimental variables for each microbiome outcome measure. Significance of each level from nested ANOVA and PERMANOVA indicated. *** *p* < 0.001.

### Precision of beta diversity within sample, subject, and cage

Given that multivariate data measurements exhibited the largest biological variability within each GM (**Supplementary Figure 3**), we next assessed the precision of this multivariate data at the technical replicate, mouse, and cage levels. We generated weighted (Bray-Curtis) and unweighted (Jaccard) distance matrices, then averaged the distances for each replicate, mouse, and cage. As expected, based on previous analyses, technical replicates exhibited the lowest dissimilarity (i.e., greatest precision) using both weighted (0.056 ± 0.005) and unweighted (0.106 ± 0.010) distance metrics (**Figure 3**). Intra-mouse dissimilarity was greater by nearly 50% (Bray-Curtis = 0.087 ± 0.015, Jaccard = 0.158 ± 0.024), whereas intra-cage dissimilarity was greater by nearly four-fold (Bray-Curtis = 0.192 ± 0.045, Jaccard = 0.307 ± 0.052). Dissimilarity was the lowest at the replicate level and progressively greater at the mouse and cage levels within both GM_Low_ and GM_High_ standardized GMs. Interestingly, we observed greater dissimilarity at each level when using unweighted, Jaccard distances (**Figure 3**, **Supplementary Figure 4**) again suggesting that – like with univariate alpha diversity measurements – presence-based measurements are most susceptible to biological and technical variability.

**Figure 3.**
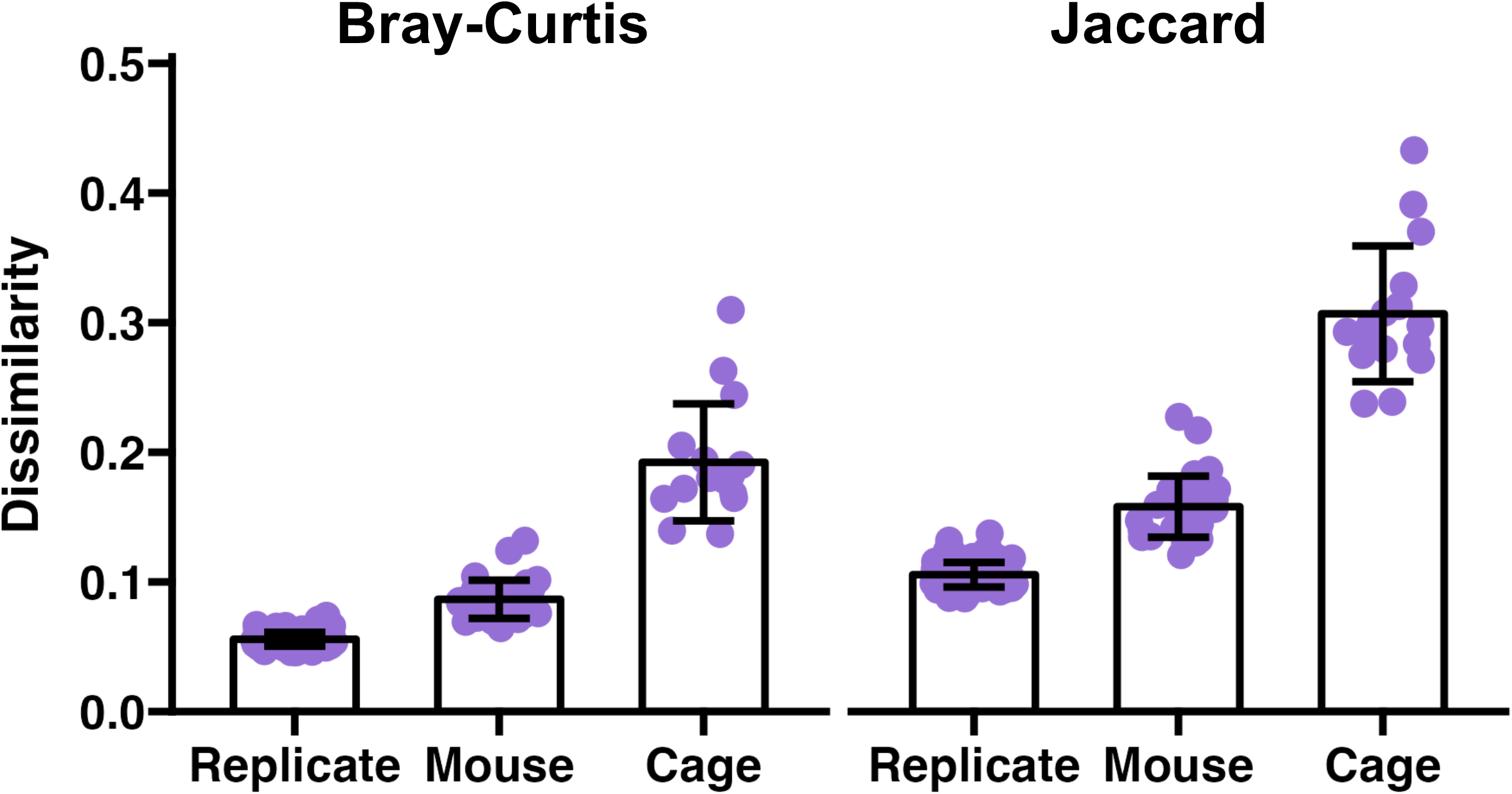
Quantifying intrasample, -subject, and -cage beta diversity. Dot plots depicting determined dissimilarity of community composition at the sample, subject, and cage levels using weighted *(left)* and unweighted *(right)* distance metrics across two standardized complex GMs.

### Repeated sampling of individual mice marginally improves effect size but markedly increases sequencing costs

Having evaluated the effects of technical and biological variability on microbiome outcome measures, we lastly determined whether taking multiple measurements (i.e., collecting multiple fecal samples) from each mouse in a study could increase effect size, thereby decreasing the number of required animals required to achieve statistical power (β = 0.8). This is based on the statistical practice of increasing the number of samples measured per mouse to decrease the variation contributed by an individual subject. To demonstrate the practical utility of these data and provide a cost analysis of the sampling strategy, we explored a scenario in which we seek to detect differences in Chao1 Index between two groups (e.g., two standardized complex GMs, drug vs. control, etc.).

Based on the present data from two standardized, complex GMs, we calculated the overall variation of the Chao1 Index at the technical replicate 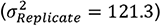, fecal sample 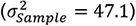, and mouse 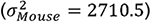 levels. Given these values, we can determine the expected variation of a given population based on the following equation: 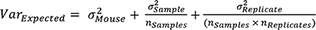[26]. Because no significant variation at the technical replicate level for Chao1 Index was observed (**Figure 2**), we can assume that 16S rRNA library preparations for each fecal sample will be performed a single time (*n_Replicate_* = 1). The estimated variation can then be used to calculate effect size between two groups using Cohen’s *d* 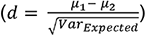. Assuming an □ of 0.05 and β of 0.8, we can determine the necessary sample size for a given difference of means and number of samples collected per mouse (*n_Samples_*).

We evaluated differences in richness (Chao1 Index) at three levels: first at the difference in means detected between GM_Low_ and GM_High_ in the present study (96) and two smaller differences (50 and 25) simulating a range of high, medium, and low mean differences, respectively. We then determined the achieved effect size (Cohen’s *d*) by increasing the number of fecal samples collected per mouse (**Figure 4**). The selected mean differences simulate medium to high effect sizes using Cohen’s *d* [27].

**Figure 4.**
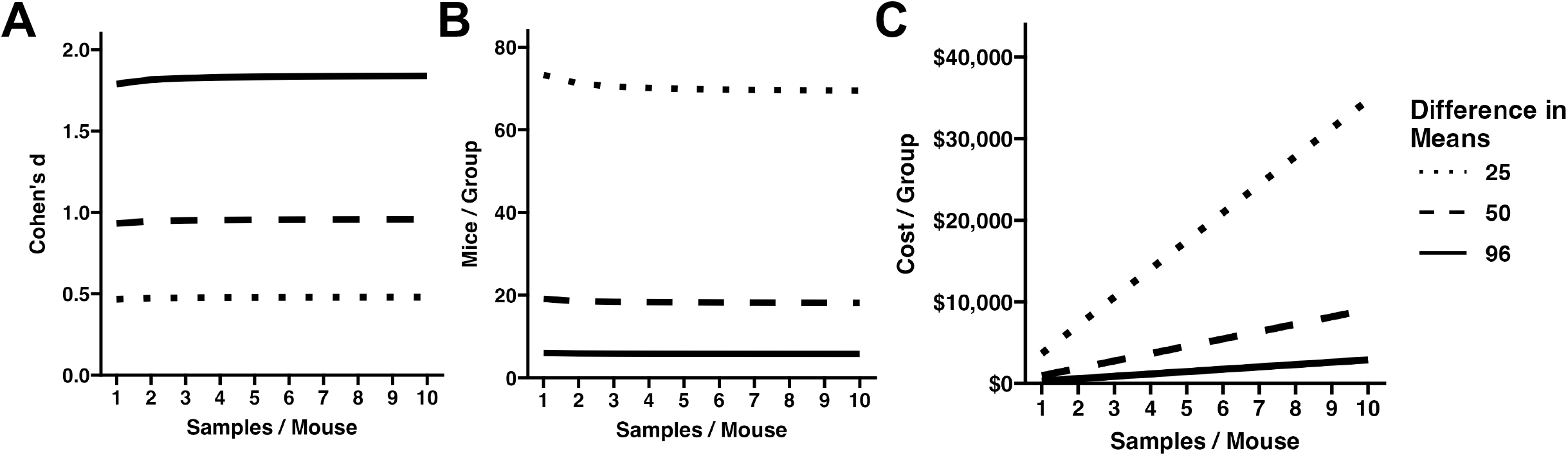
Line plots depicting the (A) achieved effect size, (B) estimated minimum sample size required per group, and (C) sequencing costs for determining varying mean differences in Chao1 Index in a two-group experimental design.

The simulated sampling of a single fecal sample per mouse achieved an effect size (Cohen’s *d*) of 1.79, 0.93, and 0.47 for the high, medium, and low mean differences, respectively (**Figure 4A**). In each case, the expected effect size was marginally increased by 1.5%, 2.0%, and 2.7% when collecting two, three, or ten samples per mouse, respectively. Assuming an alpha of 0.05 and power (β) of 0.8, we then determined the minimum number of mice per group for each simulated mean difference (**Figure 4B**). The marginal improvements to effect size by increasing the number of samples collected from each mouse translated into relatively small decreases in the minimum number of animals required per group. For example, when detecting a low mean difference, a minimum of 74 mice per group is required when collecting only one fecal sample per animal, however, collecting at least five samples per mouse decreases the minimum required sample size to 70 mice – a nearly 5% reduction. While increasing the number of samples collected from each mouse decreases the minimum number of required animals, it drastically increases sequencing costs (**Figure 4C**). Assuming library preparation and sequencing costs are $50 per sample, sequencing one sample from each mouse would cost $3,700 per group (74 mice/group × 1 sample/mouse × $50/sample), whereas, sequencing five samples from 70 mice would incur $17,500 in sequencing costs (70 mice/group × 5 samples/mouse × $50/sample).

At current sequencing costs, these data suggest that collecting and sequencing one fecal sample per mouse is the most practical sampling strategy. Based on this recommendation, we lastly calculated the minimum number of mice required per group to detect differences in common alpha diversity metrics between two groups thus providing a practical guide to sample size estimation when powering studies for microbiome alpha diversity measures (**Supplementary Figure 5**).

## Discussion

Here we have demonstrated that biological and technical variables encountered in microbiome studies contribute limited, albeit significant, variation to microbiome outcome measures in the context of an experimental variable with high intergroup variability. In our model of two supplier-origin microbiomes with robust differences in community richness and composition, we have found that the effect size of biological and technical variables is small compared to that of the experimental effect size and quantified this variation for multiple univariate alpha diversity measurements allowing for sample size calculations for future microbiome studies. The sample size estimations suggested that repeated sampling of individual mice in a two-group study design allows for marginal reductions (< 5%) in minimum sample size while considerably increasing sequencing costs. These data collectively suggest that while it is possible to reduce the number of animals required in microbiome studies by repeatedly sampling individual subjects, this strategy is currently impractical.

In using two supplier-origin microbiomes representative of Jackson Laboratory and Envigo, we captured the relative extremes of richness amongst rodent producers located in the United States [21,28]. Interestingly, in the low-richness Jackson Laboratory-origin microbiome, we observed lower precision when evaluating presence-based measurements like richness and unweighted distances. With fewer taxa present in the microbial community, subtle changes – especially in rare taxa – introduced by PCR biases or rarefaction would have a larger effect on presence-based measurements. This suggests that researchers working with lower-biodiversity microbiomes (i.e. low richness feces, skin, respiratory system, etc.) may consider taking multiple measurements of the richness of a sample(s) to determine a more precise representation of present taxa in both univariate and multivariate ecological parameters.

We then compared the relative effect size contributed by technical and biological replicates to the effect size of our experimental variable (supplier-origin GMs). Given that the effect size of technical replications was small (η^2^ = 0.027 ± 0.014) relative to the experimental variable, this suggests sequencing a fecal sample a single time is sufficient and variation introduced during 16S rRNA library preparation will likely not affect the experimental outcome. Biological variation at the mouse (η^2^ = 0.126 ± 0.086) and cage (η^2^ = 0.205 ± 0.118) levels, however, contributed appreciable effect sizes in the medium and large ranges, respectively (**Figure 2**). While these effect sizes were minimal relative to the experimental variable of supplier-origin GM (η^2^ = 0.624 ± 0.200), their contribution to variability of the dataset may be appreciable in the context of an experimental design in which the experimental variable effect size is smaller. In such case, repeated sampling of individual mice may be utilized to minimize intrasubject variation maximizing statistical power. There is potential for this strategy to be used to mitigate cage effects on microbiome measurements either alone or in combination with other strategies like decreasing housing density [13].

Beta diversity is an ecological measurement used to evaluate the dissimilarity of two communities based only on the presence (unweighted dissimilarity) or presence and abundance (weighted dissimilarity) of its members. In microbiome science, biomass collected from individual test subjects (and individual samples) are often sequenced a single time before evaluating beta diversity between experimental groups. This approach overlooks the naturally occurring variation in community composition within an individual sample or subject. Here we measured the variability in community composition within a sample, subject, and cage and found that, as expected, precision respectively decreased (i.e., dissimilarity increased) across those factors. Intrasample variability may result from technical factors like PCR or sequencing biases, intrasubject from biogeographical variation of individual fecal boli within the hindgut [18], and random developmental factors causing intra-cage host-to-host microbiome variability. Overall, understanding the precision of these beta diversity measurements is essential when developing experimental designs. As with effect size, while the technical and biological precision is relatively high in these beta diversity measurements, they may affect study outcomes in the context of experimental factors with more subtle differences in microbial composition.

A goal of this work was to provide tangible guidance for experimental designs in microbiome research. Studies are often powered for host outcome measures (i.e., behavior or disease outcomes). Likewise, it is important to appropriately power a study for GM measurements when the GM is one of the experimental outcome measures. We elected to evaluate differences in univariate alpha diversity measurements as these provide general, highly interpretable measurements of overall microbial community dynamics as opposed to multivariate and taxonomic abundance measurements which are more specific to an experimental question and design. Given that sample size calculations are dependent of the expected variability of the dataset, we simulated scenarios in which multiple fecal samples were collected from each subject to determine whether repeated sampling reduces the minimum number of animals required to detect low, medium, or high differences in means. Our simulated data suggests that repeated sampling of individual subjects reduced the minimum number of animals required by up to 5% but the financial burden of increased sequencing costs likely prohibits the practical application of this repeated sampling strategy.

Having recommended to collect a single fecal sample per mouse, we provided guidance for sample size calculations of two-group study designs. While the range of mean differences for each alpha diversity metric is large, these may serve as a guide for researchers seeking to power their studies for microbiome questions. Of note, the lower end of each mean difference range we have provided necessitates an unfeasible minimum number of animals per group (e.g., > 10,000). Recognizing this, powering studies to identify these low mean differences in alpha diversity metrics is likely not practical. For example, the Chao1 Index we calculated for the present study was observed to have a standard deviation of 13.9 and 20.8 for GM_Low_ and GM_High_ at the mouse level, respectively. Thus, it would be costly in terms of animal numbers and financial resources to perform an experiment to detect mean differences in Chao1 in groups expected mean differences less than 50 (for Chao1 Index).

The sample size estimations we have provided above pertain to univariate alpha diversity measurements only. While differences in beta diversity are frequently quantified in microbiome studies, these measurements are dependent upon pairwise comparisons of multiple samples and are difficult to quantify without pilot data relevant to the experimental groups in question [29,30]. While statistical tools have been developed to quantify the effect size of differences in beta diversity or predict effect size based on simulated data [31,32], these tools have not yet incorporated repeated sampling strategies like we have assessed here. Because of this, we were unable to determine whether repeated sampling of individual subjects could reduce sample sizes similar to what was observed with alpha diversity indices in the present study. However, based on our measurements of beta diversity precision we hypothesize that, like alpha diversity measurements, repeated sampling would likely marginally improve statistical power.

## Conclusions

Biological (i.e., intra-colony and intra-subject) and technical variability (i.e., 16s rRNA library preparation) contribute significant variation to the data in both alpha and beta diversity measurements, but the relative effect size of such variables is minimal in an experimental paradigm with large inter-group variability. Therefore, such variables should be considered in experimental designs in which the anticipated effect size of the experimental variable is significantly smaller. While repeated sampling of individual subjects reduces this variability, this approach comes with an untenable financial burden. These data highlight the value of experimental designs and practices to mitigate effects of cage and subject in microbiome research including reduced numbers of mice per cage, and intentional homogenization via shared soiled bedding.

## Abbreviations

ANOVA: Analysis of variance
B6: C57BL/6J
CV: Coefficient of variance
GM: Gut microbiome
GM_Low_: The Jackson Laboratory-origin microbiome
GM_High_: Envigo-origin microbiome
MMRRC: Mutant Mouse Resource & Research Center
NGS: Next-Generation Sequencing
PCoA: Principal Coordinate Analysis
PERMANOVA: Permutational multivariate analysis of variance

## Consent for publication

Not applicable.

## Availability of data and material

16S rRNA sequencing data are available at the Sequence Read Archive under the BioProject number PRJNA1083462. All code can be accessed at https://github.com/ericsson-lab/intrafecal_variation.

## Authors’ contributions

ZM, KG, and AE conceptualized and designed the study. ZM and KG collected samples and performed DNA extractions. ZM, KG, and AE analyzed the data and drafted the manuscript. All authors read and improved the final manuscript.

## Acknowledgements

The authors would like to thank the MU MMRRC for generously providing the CD-1 mice.

---

**Supplementary Figure 1.**
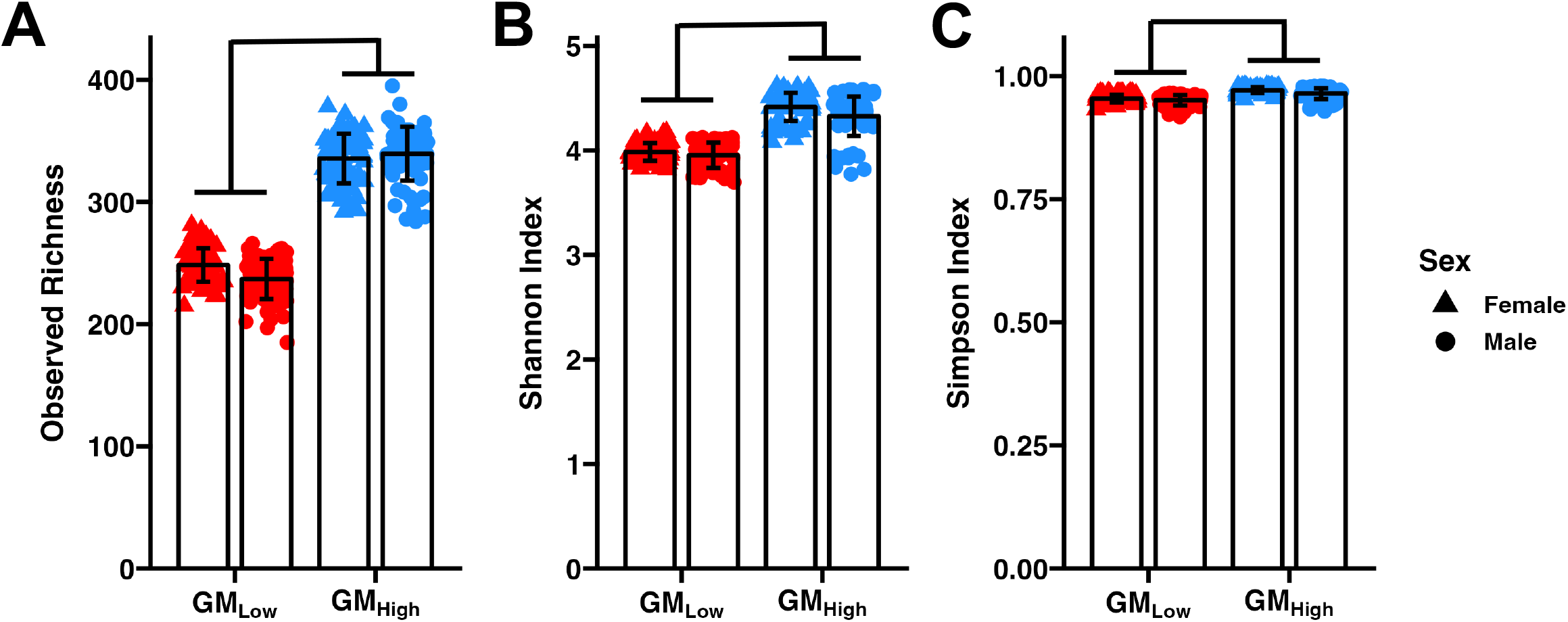
Vendor-origin microbiomes differ in richness and diversity. GM_Low_ and GM_High_ microbiomes differ in (A) observed richness (GM: F = 289, *p* < 0.001), Shannon Index (GM: F = 67.5, *p <* 0.001), and Simpson Index (GM: F = 23.5, *p <* 0.001), two-way ANOVA.

**Supplementary Figure 2.**
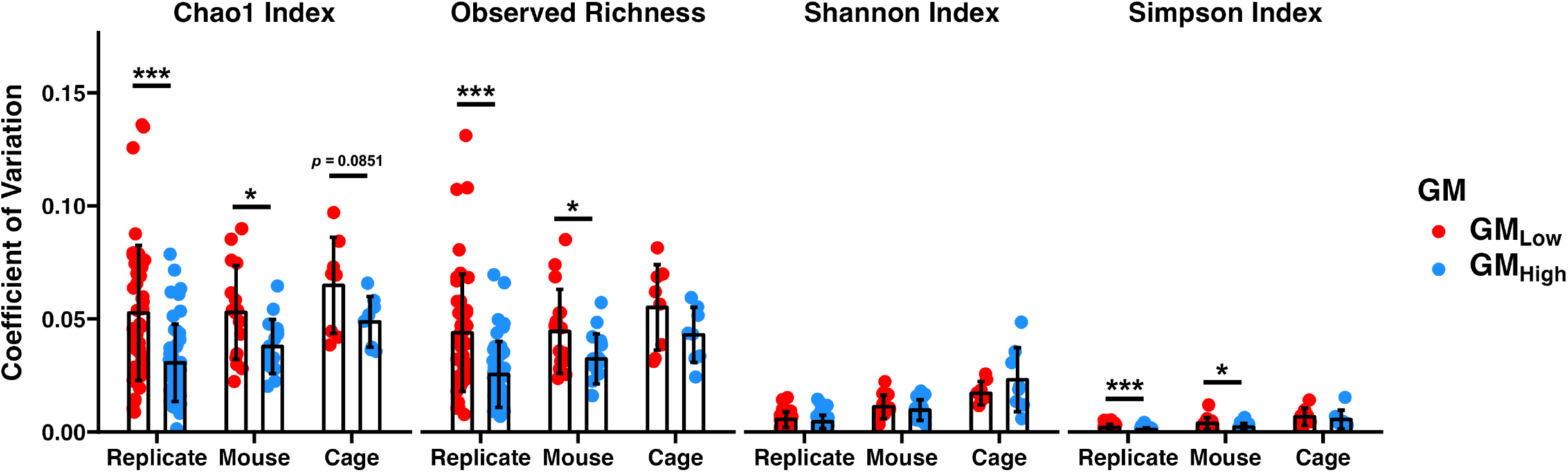
Low-richness GM exhibits lower precision in presence-based alpha diversity measurements. Dot plots depicting coefficient of variation measured at the replicate, mouse, and cage levels in GM_Low_ and GM_High_ for Chao1 Index, Observed Richness, Shannon Index, and Simpson Index (*left to right*). \**p*< 0.05, \*\*\**p*< 0.001, t test between GMs for indicated level.

**Supplemental Figure 3.**
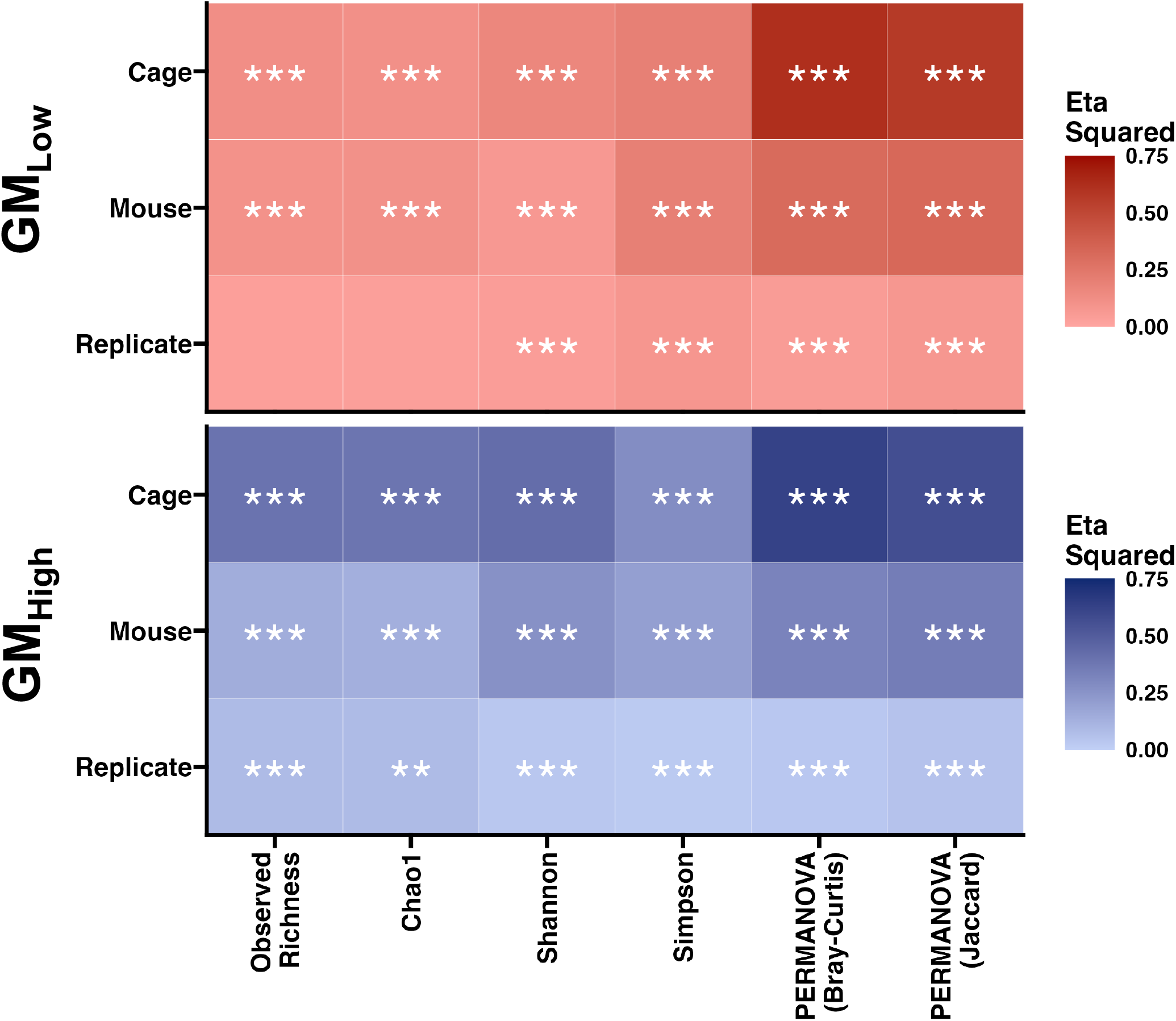
Evaluating the relative effect size of biological and technical variables within standardized complex GMs. Heatmaps depict the relative effect size contributed by technical and biological variables in GM_Low_ *(top)* and GM_High_ *(bottom)* for each microbiome outcome measure. Significance of each level from nested ANOVA and PERMANOVA indicated. *** *p* < 0.001.

**Supplementary Figure 4.**
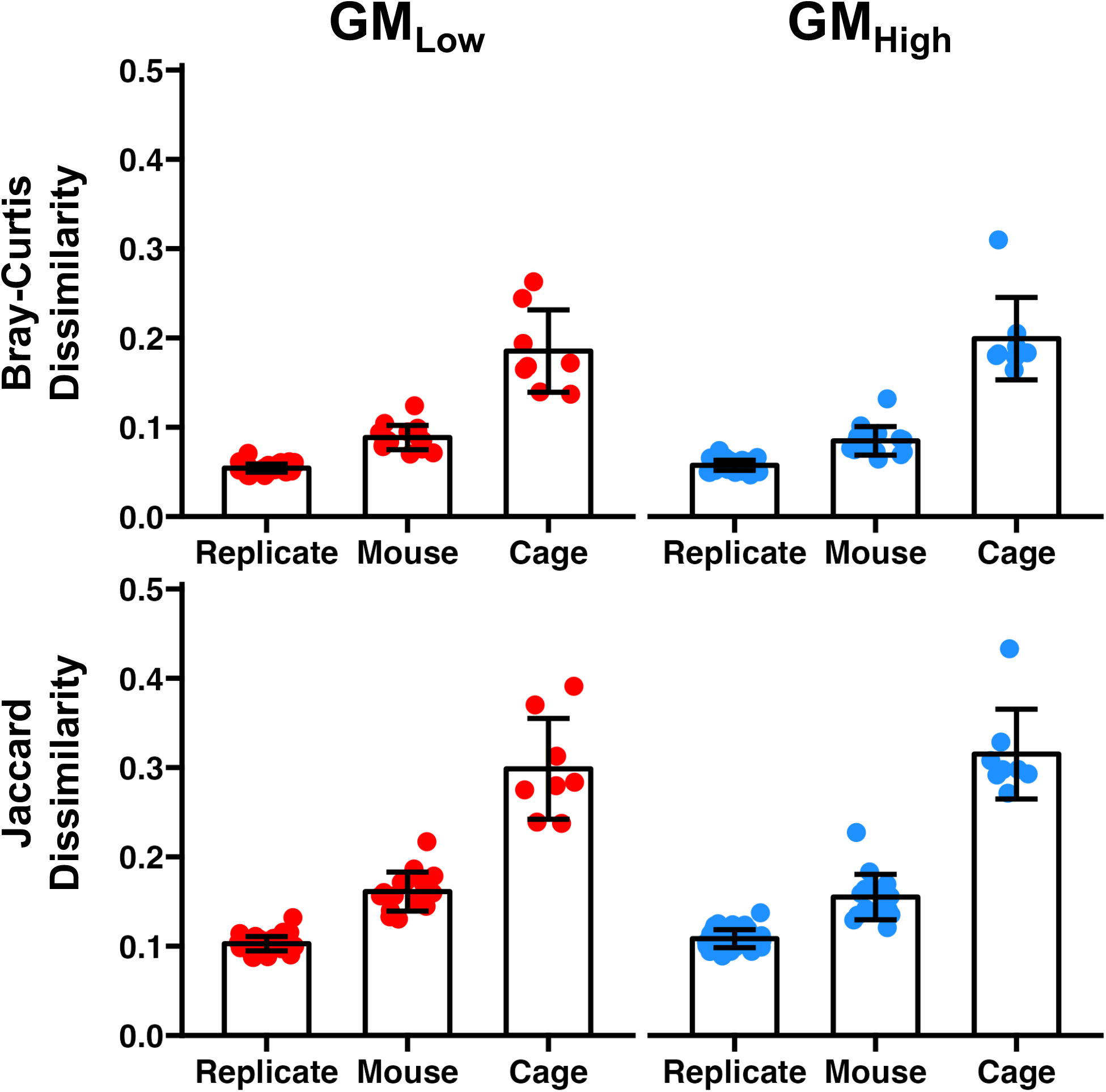
Quantifying intrasample, -subject, and –cage beta diversity within standardized complex GMs. Dot plots depicting determined dissimilarity of community composition at the sample, subject, and cage levels using weighted *(top)* and unweighted *(bottom)* distance metrics within GM_Low_ *(left)* and GM_High_ *(right)*.

**Supplementary Figure 5.**
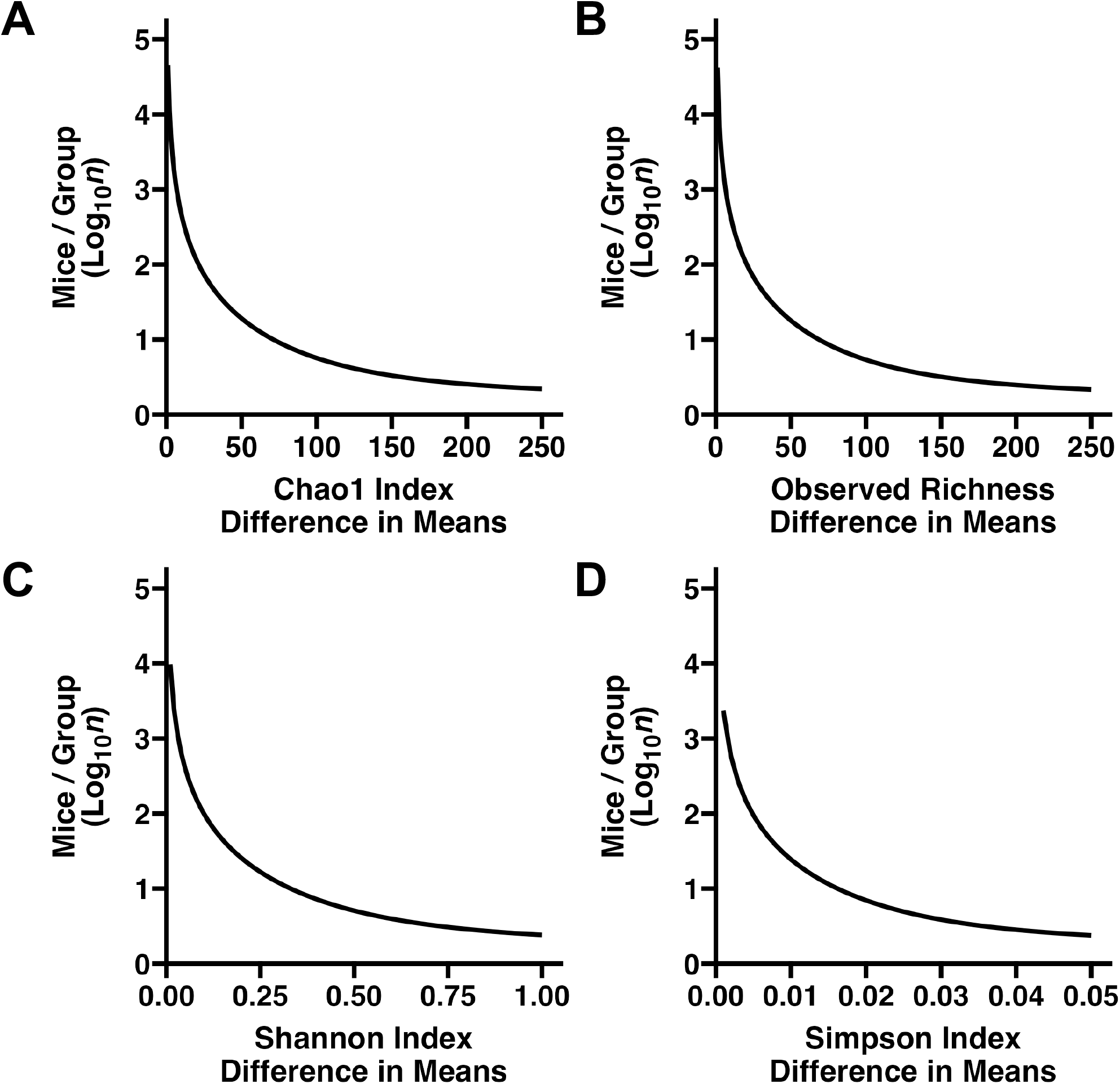
Line plots depicting the minimum number of mice per group required to detect varying mean differences in **(A-B)** richness and **(C-D)** alpha diversity when collecting and sequencing one sample per mouse.

